# A Population-Specific PARP1 Gene Variation Modulates PARP Trapping

**DOI:** 10.1101/2025.11.11.687916

**Authors:** Jin Cai, Ramya Billur, Michael S. Cohen, Andreas G. Ladurner, Ben E. Black

**Author notes:** These authors contributed equally (Jin Cai, Ramya Billur). Correspondence (A.G.L.), (B.E.B.).

## Abstract

Poly-(ADP-ribose) polymerase inhibitors (PARPi) block NAD^+^-binding pocket of PARP1, inhibiting PAR synthesis. However, they differ in their ability to retain PARP1 on damaged DNA to induce synthetic lethality in homologous recombination-deficient (HRD) cancer. Allosteric enzymatic activation requires destabilization of the helical domain (HD) of PARP1 and is indispensable for activation and chromatin retention induced by distinct PARPi. Here we report that the effect of the clinical PARPi talazoparib is robustly impacted by a common human polymorphism within the HD. PARP1^V762^ greatly enhanced talazoparib-driven allosteric retention on chromatin, prolonged XRCC1 recruitment, and enhanced cell killing. Talazoparib switches from Type-II PARPi behavior in PARP1^A762^ to allosteric, pro-retention Type-I behavior for PARP1^V762^. Thus, both PARPi efficacy and dose-limiting tolerability depends on PARP1 allele, motivating variant-guided cancer therapies.

**TEASER:** One of the four FDA-approved PARPi drugs, talazoparib, is modulated by a PARP1 SNP that is widespread in the population.

## INTRODUCTION

Poly-(ADP-ribose) polymerase inhibitors (PARPi) are the first clinically approved drugs that exploit the established genetic concept of synthetic lethality^1,2^. Their main target is the human nuclear enzyme PARP1 which contribute to ∼90% of cellular poly-ADP-ribosylation (PARylation) activity^3^. PARP1 undergoes an allosteric transition and enzymatic activation upon recognizing and binding to DNA lesions. This is mediated by communication between its DNA binding domain and auto-inhibitory helical domain (HD). Upon DNA binding, the HD helices partly unfold, which allows the cofactor NAD^+^ to bind the catalytic domain^4–6^. PARP1 then synthesizes poly-(ADP-ribose) chains (PAR) from NAD^+^ to modify itself and other substrates. In turn, this promotes the self-dissociation of PARP1 from chromatin and promotes the recruitment of DNA repair machinery, including chromatin remodelers, to the lesions^7–10^.

Distinct PARPi have been developed over the last 20 years for use in the cancer clinic. Their shared function reflects the orthosteric inhibition of catalytic PARP1 activity, so all PARPi directly compete with the cofactor NAD^+^ for binding to the enzyme’s catalytic domain, shutting down PAR synthesis and in several cases also promoting the physical retention of PARP1 enzyme on chromatin^11^. Enhanced PARP retention, often referred to as “PARP trapping”, indicates the increased association of PARP1 enzyme to chromatin in the presence of a PARPi. PARP trapping blocks the progression of DNA replication forks, inhibits fork reversal and restart, and requires homologous-recombination-mediated DNA repair (HR) for resolution. In turn, such trapping is thought to contribute to the synthetic lethality in cancers deficient in HR repair (HRD), including a significant fraction of ovarian cancer^1,2^,^12^. The PARPi olaparib, rucaparib, niraparib and talazoparib have all been approved by the FDA and EMA, being employed in the clinic primarily as maintenance therapy for recurrent breast, ovarian, prostate and pancreatic cancer^13,14^. Despite their shared mechanism of catalytic inhibition, these PARPi differ markedly in their *in vitro* and cellular potency, efficacy, dose-limiting toxicity, and notably also their ability to trap PARP enzymes on chromatin, factors that are partly interrelated, but not yet fully understood mechanistically. In particular, how PARP trapping contributes to therapeutic benefit while also impacting safety across different cancer and patient contexts remains incompletely understood.

Indeed, while PARPi endow superior efficacy with progression-free survival benefits over standard-therapy group or placebo^15–24^, PARPi use also is associated with dose-limiting toxicities, such as myelosuppression, gastrointestinal disorders and fatigue that often require dose reductions or the interruption of therapy. Grade 3 (or greater) anemia occurred in 20% to 40% of clinical study participants^18^, as well as high occurrence of grade ≥3 neutropenia and thrombocytopenia, plus rare cases of myelodysplastic syndrome (MDS), AML and pneumonitis. Coupled to widespread PARPi resistance in settings where patients initially respond well to PARPi^25^, the field is exploring alternatives, such as the development of improved PARP1-selective inhibitors, or targeting orthogonal DNA damage and repair (DDR) factors, such as the PAR glycohydrolase PARG^26^,the ubiquitin-specific protease USP1 in replication stress^27^ or the allosterically-regulated, PAR-dependent remodeler ALC1/CHD1L^28^.

Beyond PARPi’s inhibition of PARylation activity, several PARPi promote the physical retention of PARP1 on chromatin. Based on the reverse allostery induced by individual PARPi from the HD subdomain (of the catalytic domain) to DNA binding domains, PARPi are classified into three types: Type-I PARPi contact the αF helix of PARP1, initiate reverse allostery and thus enhance interdomain interactions and DNA-binding capacity; to date, no clinical PARPi are known to act through this mechanism. Type-II PARPi bind to PARP1, but induce minor or negligible changes in interdomain interactions, resulting in modest alterations to DNA binding, including the clinical-relevant PARPi talzoparib and olaparib. In contrast, Type-III PARPi disrupt interdomain interactions to reduce the affinity of PARP1 for DNA^29,30^. The interaction with the αF helix in the HD subdomain of PARP1 and resulting changes in backbone dynamics have emerged as critical in determining chromatin retention and reverse allostery, influencing its capacity to interfere with DNA replication during PARPi treatment.

Further, our insights into how single nucleotide polymorphisms (SNPs) in the PARP1 gene impact the function of therapeutic PARPi are limited. Considering the increasing use of PARPi in the cancer clinic, we thus sought to investigate this question. Interestingly, a common human polymorphism impacts codon 762, located within the regulatory αF helix on the HD domain of PARP1, which results in the substitution of a valine with an alanine residue^31^. PARP1^A762^ is the minor variant overall, with an occurrence in around 45% of persons with Asian ancestry, 15% of Caucasians and almost absent in persons with African ancestry^32^

(approximately 5%). Notably, almost all biochemical and drug discovery research with PARP1 has been conducted with the minor PARP1^A762^ variant over the last ∼35 years and not with the PARP1^V762^ variant, whose frequency in some human populations is almost one hundred percent. It is fair to assume that most of our knowledge on how distinct PARPi impact PARP1 activity, biochemical function and therapeutics response has been gathered from studying the less frequent/rare human PARP1 variant.

Considering that variant-specific responses may impact the clinical benefit vs. tolerability balance associated with the approved, non-selective PARPi, we took a closer look at how PARPi impact the two predominant PARP1 variants present in the human population. Interestingly, it has been proposed that PARP1^A762^ may confer greater cancer predisposition^33–36^. At the biochemical level, PARP1^A762^ exhibits lower enzymatic activity than PARP1^V762^, with a reported mild decrease of PAR synthesis *in vitro*^37,38^ and less PAR formation upon hydrogen peroxide induction *in vivo*^39^, for example. However, nothing is known as to whether and how the PARP1 variants differ in their response to PARPi, in particular we do not know whether and how they impact PARPi-induced PARP1 retention, which is thought to represent the key feature of PARPi that is most closely linked to the efficacy and safety profile of PARPi therapies. Importantly, V/A762 lie in the regulatory, HD domain helix αF, a region critical for allosteric PARP1 activation. A population-specific PARP1 gene variation thus situates to the edge of the canonical PARPi binding site within the PARP1 active site (**Figure S1**) and has been shown to be in direct contact with distinct PARPi.

Here we sought to test whether and how the two human PARP1 SNP variants differ in their response to clinical PARPi. We used imaging, cell biology and biophysical assays to probe how distinct PARPi, including novel PARP1-selective inhibitors, impact the structure and function of PARP1. We tested for the impact of PARP1 variants on cellular viability in the presence of distinct PARPi and on the recruitment of cellular factors responding to DNA damage. Our results show that PARP1^V762^ was retained stronger on chromatin compared to PARP1^A762^ when inhibited by talazoparib. The change of valine 762 to alanine converted talazoparib from a Type-II PARPi to a Type-I PARPi. Further, HXMS analysis uncovered how activated PARP1^V762^, in the presence of talazoparib, remarkably unfolded the HD subdomain while reinforcing interdomain interactions. In agreement with the stronger retention of PARP1 on damaged chromatin, PARP1 V762-bearing cells were associated with persistent DNA damage and higher sensitivity towards talazoparib treatment. Our results suggest that efficacy, toxicity and other clinical parameters may be impacted by this widespread PARP1’s gene variation. Our data provide insights on why some patients may respond differently to the therapeutic use of PARPi when targeting HR-deficient tumors.

## RESULTS

### Talazoparib induces a higher retention of PARP1^V762^ at irradiated sites

To investigate the impact of PARP1 variants on PARP1 functionality and PARPi efficacy, we monitored the retention kinetics of the two variants at DNA damage sites under PARPi treatment using live-cell imaging following micro-irradiation. U2OS PARP1-knockout (ΔPARP1) cells stably expressing GFP-tagged PARP1^V762^ or PARP1^A762^ (**Figure S2A**), were exposed to DNA damage using 355 nm laser microirradiation and the recruitment of the GFP-tagged PARP1 variants to the DNA lesions was recorded and quantified over 15 minutes (**Figure 1A**). In the control group, there was a rapid and dynamic recruitment of PARP1 as previously reported^40^, peaking at around 1 minute post irradiation, and dissociating gradually afterwards, with no notable difference between the two variants (**Figure 1B**).

**Figure 1.**
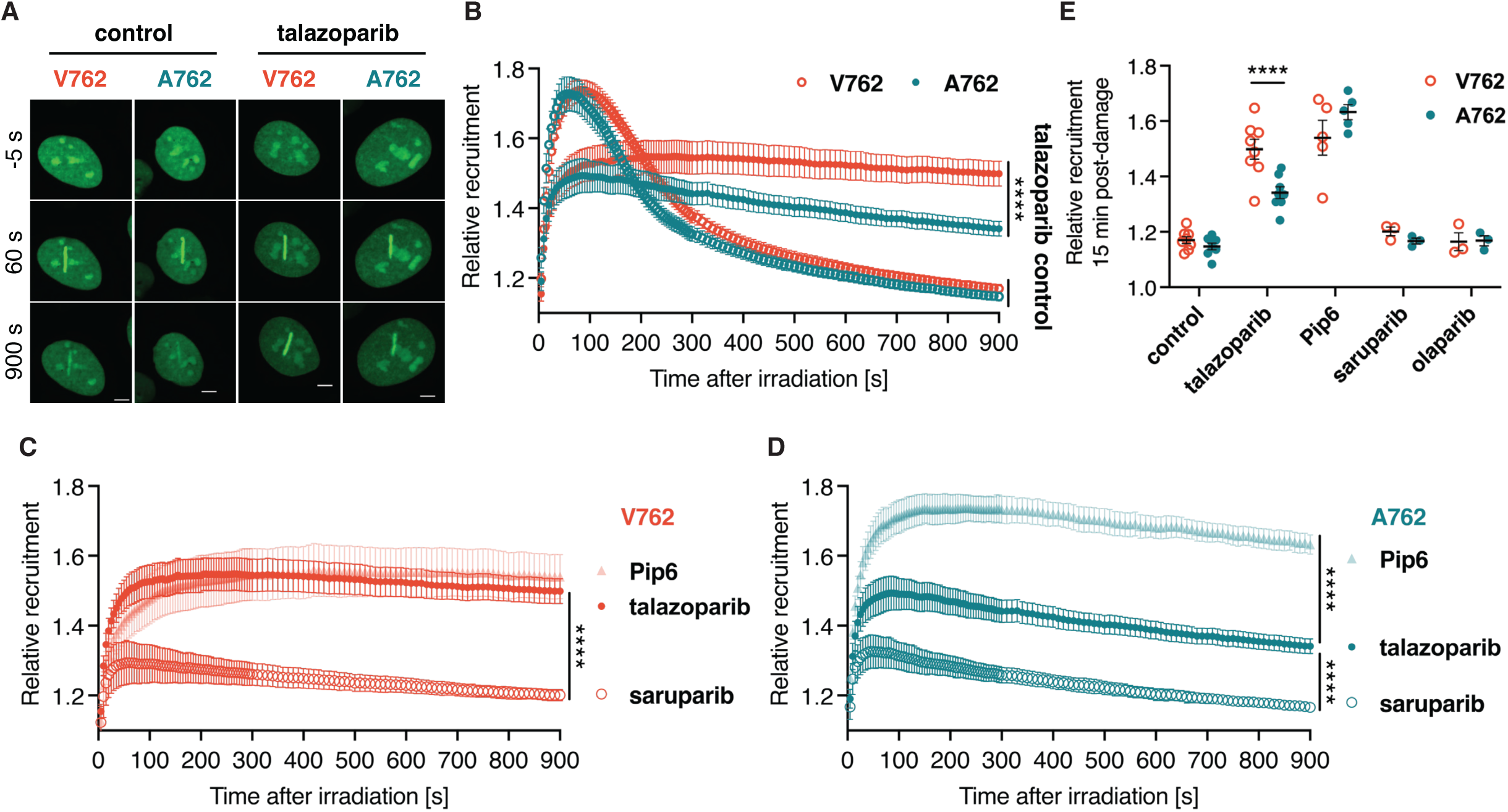
Talazoparib induces higher retention of PARP1^V762^ at irradiated sites. (**A**) Representative images of recruitment of GFP-tagged PARP1^V762^ or PARP1^A762^ to micro-irradiated sites before or 60 seconds and 900 seconds after micro-irradiation in U2OS ΔPARP1 cells stably expressing PARP1-GFP V/A762. Cells were treated with or without 1 μM of talazoparib before irradiation. Scale bar 5 μm. (**B**) Recruitment kinetics of GFP-tagged PARP1^V762^ or PARP1^A762^ to micro-irradiated sites over 15 minutes in U2OS ΔPARP1 cells stably expressing PARP1-GFP V/A762. Cells were treated with or without 1 μM of talazoparib for 1 hour before irradiation. Mean ± s.e.m. of n = 8 independent experiments with Student’s t test. (**C–D**) Recruitment kinetics of GFP-tagged PARP1^V762^ (C) or PARP1^A762^ (D) to micro-irradiated sites over 15 minutes in U2OS ΔPARP1 cells stably expressing PARP1-GFP V/A762. Cells were treated with or without 1 μM of indicated PARPi for 1 hour before irradiation. Mean ± s.e.m. of n = 8 (talazoparib), 5 (Pip6) or 3 (saruparib) independent experiments with Student’s t test. (**E**) Relative recruitment of GFP-tagged PARP1^V762^ or PARP1^A762^ at micro-irradiated sites 15 minutes post-damage from (B–D). Cells were treated with or without 1 μM of indicated PARPi for 1 hour before irradiation. Mean ± s.e.m. of n = 8 (control and talazoparib), 5 (Pip6) or 3 (saruparib and olaparib) independent experiments with Student’s t test.

Upon a short exposure to talazoparib, however, we observed reduced recruitment and a long-lasting retention of PARP1 at the induced DNA damage sites. Strikingly, the retention of PARP1^V762^ was significantly higher than that of PARP1^A762^. We also tested the retention of PARP1 variants with the treatment of a preclinical Type-I PARPi, Pip6^41^, together with a distinct Type-II PARPi, olaparib. For PARP1^V762^, both talazoparib and Pip6 induced extensive PARP1 retention in the imaging assay, implying that talazoparib acts as Type-I PARPi on PARP1^V762^, the same as Pip6 (**Figure 1C**).

For PARP1^A762^, however, only Pip6 triggered an intensive retention of the PARP1 variant, while talazoparib retained it mildly, in line with the previous classification of talazoparib as Type-II PARPi^29,41^, which was established with PARP1^A762^ (**Figure 1D**). In addition, olaparib and the newly-developed PARP1-selective PARPi sarubparib (AZD5305), showed a low and indistinguishable retention for either of the variants, similar to the retention in control group (**Figure 1E**). Taken together, our imaging assays show that the PARP1^V762^ and PARP1^A762^ variants differ in their recruitment and retention kinetics at induced DNA damage sites. Notably, the two human PARP1 variants differ in their behavior with the potent PARPi talazoparib.

### Talazoparib promotes reverse allostery in PARP1^V762^

To measure the impact of talazoparib-induced changes in protein backbone dynamics, we employed hydrogen/deuterium exchange assay coupled to mass spectrometry (HXMS). We used the same dumbbell model of DNA damage previously used to study PARP1^A762^ activation and PARPi-induced allosteric impact on DNA binding^5,29^. HXMS is a powerful technique that reveals insights into protein structure, dynamics, and function by measuring the exchange of backbone amide hydrogens with deuterium in solution^43^. Stable structures such as α-helices involving amide hydrogens are protected from HX. However, when local transient unfolding and refolding occurs, the amide hydrogen can undergo exchange with deuterium from D_2_O. This provides a window into conformational dynamics of the enzyme’s structure. We first found that both PARP1^A762^ and PARP1^V762^ have measurable differences in DNA damaged-induced communication that results in rapid unfolding/refolding of the HD (**Figures S3A-S3C**). This is consistent with difference between the two versions subtly impacting enzymatic activity *in vitro*^37–39^. However, how distinct PARPi interact with the distinct PARP1 isoforms was not known. Fortunately, the number of available PARP1 structures bound to several clinical and experimental PARPi has recently increased, allowing us to pinpoint any potential differences in the interaction of the PARPi with PARP1^V/A762^. Among those, 4PJT (PARP1^V762^) and 4UND (PARP1^A762^) represent crystal structures where the catalytic domain (CAT) was bound to talazoparib^44,45^. Interestingly, in the PARP1^A762^ structure, the contacts between fluorophenyl and triazol rings of talazoparib and E763 and D766 of the αF helix were altered compared to in the PARP1^V762^ structure (**Figure S1**). Prompted by this observation, we investigated whether talazoparib binding to PARP1^V762^ would induce distinct allosteric changes within the PARP1 enzyme compared to the PARP1^A762^ variant.

Given that the 100 s timepoint was the most informative in our previous HXMS studies^29,46^, we performed our HXMS to measure the impact of talazoparib binding to PARP1^V762^ already engaged to SSB DNA. Our data revealed that the peptides in αB and the C-terminal of αF showed increased exchange. This suggests that the binding of talazoparib promoted HD unfolding (**Figures 2A, 2C-2E** and **S4**). These are the same helices that were unfolded during DNA-mediated PARP1 activation. Interestingly, talazoparib reduced exchange at αE, likely due to enhanced interactions between WGR and αE. Increased protection in Zn3 and WGR domains was also observed, suggesting stabilization of multi-domain interfaces (**Figures 2A, 2C-2E** and **S4**). To further probe allosteric effects of the PARPi in the PARP1^V762^ variant, we tested whether olaparib – a Type-II PARPi for PARP1^A762^ known to sterically clash with the central region of αF, – could induce reverse allostery in PARP1^V762^ engaged to SSB DNA. Our data indicated that olaparib did not change any dynamics in HD peptides or in the inter-domain contacts and DNA binding domains, with the only significant changes confined to peptides within the ART active site, consistent with stable olaparib engagement (**Figures 2B, 2C, 2F** and **S4**). Together, our results indicate that talazoparib rapidly sampled the PARP1^V762^ HD to unfolded state and further stabilized the multidomain interfaces necessary to allosterically trap PARP1^V762^ onto a broken DNA. In contrast, the PARP1^V/A762^ variants showed essentially similar behavior with the PARPi olaparib.

**Figure 2.**
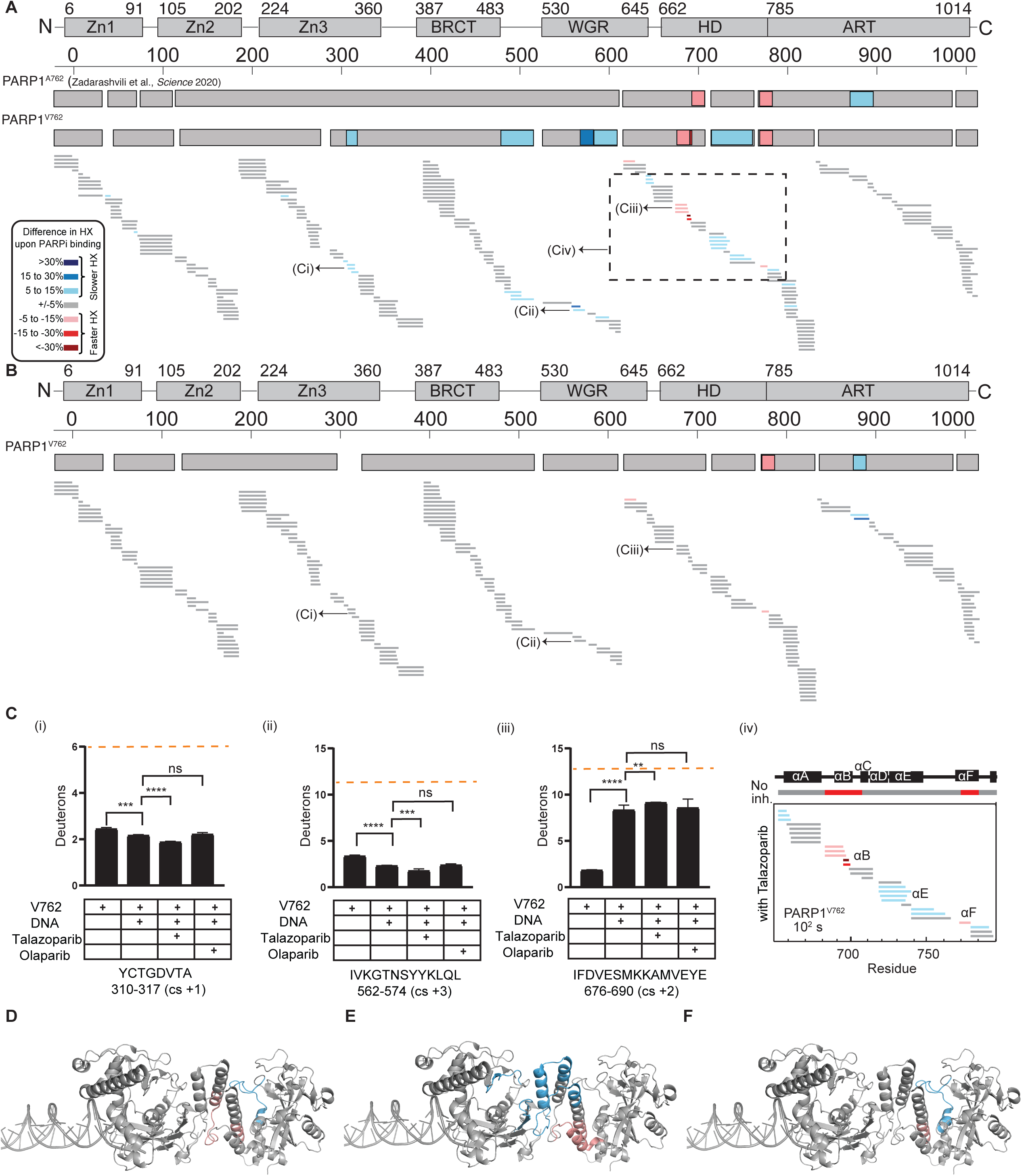
Talazoparib promotes reverse allostery in PARP1^V762^. (**A**) The percentage difference in deuteration between PARP1^V762^+DNA complex and PARP1^V762^ +DNA+talazoparib complex at 100 s was shown as a HXMS difference plot. Every horizontal bar represents a peptide. The peptides that are in gray represent no change in percentage difference. Peptides in blue represent increased protection and the peptides in red represent decreased protection. The consensus HX difference is shown on the top. The white gaps indicate no peptide coverage. (**B**) The percentage difference in deuteration between PARP1^V762^+DNA complex and PARP1^V762^ +DNA+olaparib complex at 100 s is shown as a HXMS difference plot. Every horizontal bar represents a peptide. The consensus HX difference is shown on the top. The white gaps indicate no peptide coverage. Color key for binning of HX difference is shown at the left corner. (**C**) HX of representative peptides from Zn3, WGR, and HD (i–iii) for V762, V762+DNA, V762+DNA+talazoparib, V762+DNA+olaparib complexes are shown here. An average from three replicates with error bars representing SD and asterisks indicating *P* < 0.05 from two-sided t-test between any two protein states is shown. Maximum number of deuterons (maxD, which is the total number of residues minus the first two residues due to severe back-exchange within the experimental timescale and minus the number of prolines due to no backbone amide hydrogen) exchanged by the representative peptides is indicated by the orange dashed line. (iv) percentage difference between PARP1^V762^+DNA complex and PARP1^V762^ +DNA+talazoparib complex in the HD region of PARP1. Peptides are shown as I panel A. (**D**) Consensus HXMS percentage differences between PARP1^A762^+DNA and PARP1^A762^+DNA+talazoparib complex mapped on the crystal structure of PARP1 on SSB (PDB 4DQY). (**E**) Consensus HXMS percentage differences between PARP1^V762^+DNA and PARP1^V762^+DNA+talazoparib complex mapped on the crystal structure of PARP1 on SSB (PDB: 4DQY). (**F**) Consensus HXMS percentage differences between PARP1^V762^+DNA and PARP1^V762^+DNA+olaparib complex mapped on the crystal structure of PARP1 on SSB (PDB: 4DQY).

### Talazoparib promotes higher retention of PARP1^V762^ at methylated DNA lesions

Our imaging and biophysical assays revealed that the PARPi talazoparib selectively impacts the PARP1^V762^ variant in a manner that may lead to higher PARP trapping. We thus employed an orthologous method in the field to directly probe the retention (trapping) of PARP1 at methylated DNA lesions. We employed an immunofluorescence-based PARP trapping assay^47^ and found that talazoparib distinctly impacts PARP1 trapping of the PARP1^V762^ and PARP1^A762^ variants. U2OS cells expressing the two distinct GFP-PARP1 variants were pre-extracted to remove soluble PARP1 proteins and the chromatin fraction was then fixed and imaged for PARP1-GFP signal (**Figure 3A**).

**Figure 3.**
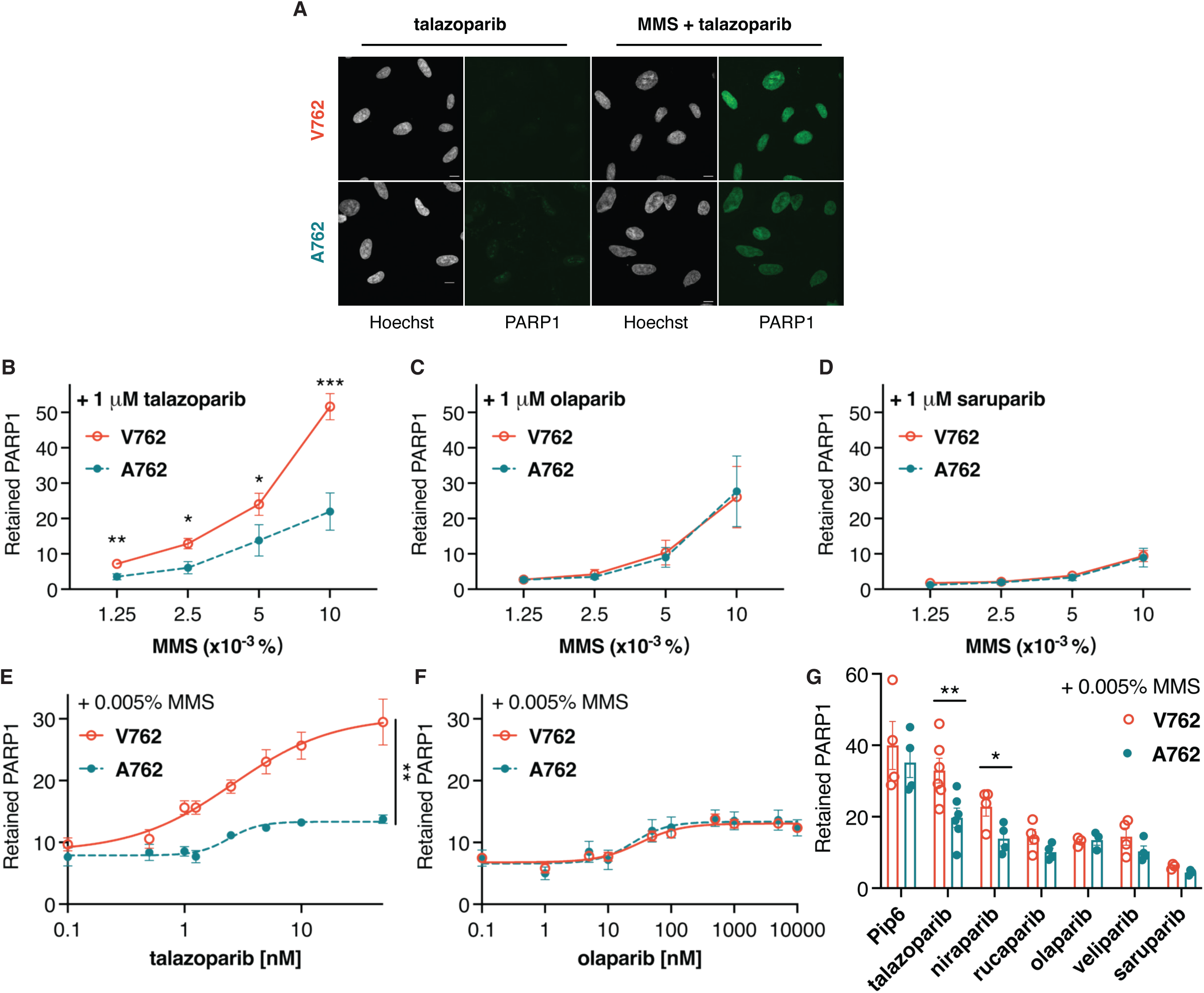
Talazoparib promotes higher retention of PARP1^V762^ also at methylated lesions. (A) Representative images of chromatin retention of GFP-tagged PARP1^V762^ or PARP1^A762^ by immunofluorescence in U2OS ΔPARP1 cells stably expressing PARP1-GFP V/A762. Cells were treated with 1 μM of talazoparib in the presence and absence of 0.01% MMS for 4 hours. Scale bar 10 μm. (**B–D**) Relative chromatin retention of GFP-tagged PARP1^V762^ or PARP1^A762^ by immunofluorescence in U2OS ΔPARP1 cells stably expressing PARP1-GFP V/A762. Cells were treated with MMS titration along with 1 μM of talazoparib (B), olaparib (C) and saruparib (D) for 4 hours. Mean fluorescence intensity of PARP1-GFP variants was subtracted to the background intensity and normalized to the PARPi-only group. Mean ± s.e.m. of n = 6 (B) or 4 (C-D) independent experiments with Student’s t test. (**E–F**) Relative chromatin retention of GFP-tagged PARP1^V762^ or PARP1^A762^ by immunofluorescence in U2OS ΔPARP1 cells stably expressing PARP1-GFP V/A762. Cells were treated with a titration of talazoparib (E) and olaparib (F) in the presence of 0.005% of MMS for 4 hours. Mean fluorescence intensity of PARP1-GFP variants was subtracted to the background intensity and normalized to non-treated group. Mean ± s.e.m. of n = 3 independent experiments with Student’s t test. (**G**) Relative chromatin retention of GFP-tagged PARP1^V762^ or PARP1^A762^ by immunofluorescence in U2OS ΔPARP1 cells stably expressing PARP1-GFP V/A762. Cells were treated with 0.005% of MMS along with 1 μM of different PARPi for 4 hours. Mean fluorescence intensity of PARP1-GFP variants was subtracted to the background intensity and normalized to non-treated group. Mean ± s.e.m. for talazoparib (n = 6) or other PARPi (n=4 independent experiments with Student’s t test).

Under a titration of the DNA alkylating-agent methyl methanesulfonate (MMS), to chemically induce global DNA damage for 4 hours in the presence of 1 μM of talazoparib, we observed a dose-dependent increase of PARP1 retention (**Figure 3B**). Remarkably, the chromatin retention of PARP1^V762^ was greater than that of PARP1^A762^. MMS treatment together with olaparib also induced a concentration-dependent increase of PARP1 retention, but to a lower degree, with a similar fold change to that of MMS co-treated with talazoparib on PARP1^A762^ (**Figure 3C**). Interestingly, there was no measurable difference between the two PARP1 variants when treating cells with MMS together with olaparib. This was also observed for the co-treatment of MMS and saruparib, a PARP1-selective inhibitor currently in clinical testing. Noteworthily, saruparib trapped PARP1 to a lower degree compared to that of both talazoparib and olaparib (**Figure 3D**).

In an orthogonal set of experiments, we titrated increasing concentrations of PARPi at fixed MMS concentrations, which also showed a dose-dependent increase of PARP1 retention as talazoparib concentration was increased, especially for the PARP1^V762^ variant (**Figure 3E**). An intensive entrapment of PARP1^V762^ and a remarkable difference between the variants was already seen at the low cellular concentration of 1 nM of talazoparib for the PARP1^V762^ variant. In contrast, olaparib titration revealed no differences in PARP1 trapping under the utilized conditions, a much lower retention and no visible difference between the variants observed at micromolar concentrations of olaparib (**Figure 3F**). We further tested all the clinical-relevant PARPi at the concentration of 1 μM in the presence of low dosage of MMS (**Figure 3G**). The ability of PARP1 retention increased from veliparib to talazoparib as reported^11^. More importantly, there was only a retention difference between the variants upon treatment of talazoparib and niraparib. These data reveal that a coding SNP at residue 762 of PARP1 within human populations robustly impacts the trapping of PARP1 induced by certain PARPi.

### PARP1^V762^ confers hypersensitivity toward the PARPi talazoparib

Considering the difference on the retention of PARP1 variants with the PARPi talazoparib, we next explored whether PARP1 variants impacted the effectiveness of DNA repair. Along with the retained-PARP1 analysis via the immunofluorescence-based trapping assay, we thus quantified the intensity of γH2AX, the phosphorylated histone H2A variant, which assists in relocating the DNA repair machinery to lesions and is commonly exploited as a sensitive DNA damage marker^48^. Under the incubation of 0.01% of MMS in the presence of 1 μM talazoparib, we observed a mild increase of pan-nuclear γH2AX signal in PARP1^V762^ bearing cells, implying more DNA damage in PARP1^V762^ bearing cells (**Figure 4A**).

**Figure 4.**
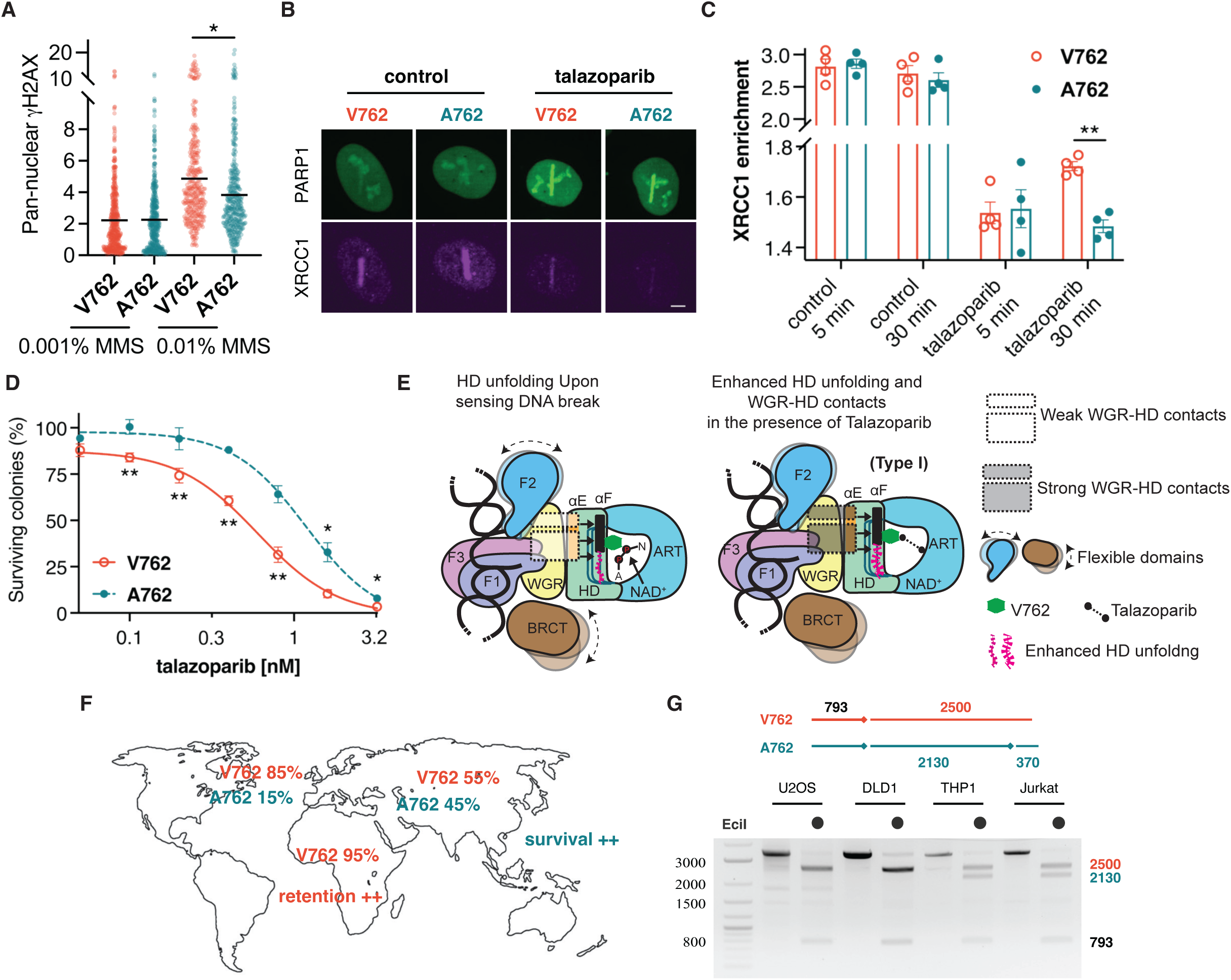
PARP1^V762^ confers hypersensitivity toward talazoparib. (A) Relative γH2AX levels by immunofluorescence in U2OS ΔPARP1 cells stably expressing GFP-tagged PARP1^V762^ or PARP1^A762^. Cells were treated with 0.001% or 0.01% of MMS in the presence of 1 μM of talazoparib for 4 hours. Mean fluorescence intensity of γH2AX was subtracted to background intensity and normalized to the PARPi-only group. Mean ± s.e.m. of n = 6 independent experiments with Student’s t test. Representative images of recruitment of GFP-tagged PARP1^V762^ or PARP1^A762^ and XRCC1 by immunofluorescence after micro-irradiation in U2OS ΔPARP1 cells stably expressing GFP-tagged PARP1^V762^ or PARP1^A762^. Cells were treated with or without 1 μM of talazoparib before irradiation and fixed 30 minutes post irradiation. Scale bar 5 μm. (**C**) Relative recruitment of XRCC1 by immunofluorescence after micro-irradiation in U2OS ΔPARP1 cells stably expressing GFP-tagged PARP1^V762^ or PARP1^A762^. Cells were treated with or without 1 μM of talazoparib before irradiation and fixed 30 minutes post irradiation. Mean fluorescence intensity of XRCC1 at micro-irradiated sites was subtracted to the background intensity and normalized to the nuclear signal. Mean ± s.e.m. of n = 4 independent experiments with Student’s t test. (**D**) Clonogenic survival of U2OS ΔPARP1 cells stably expressing GFP-tagged PARP1^V762^ or PARP1^A762^. Cells were incubated with titration of talazoparib for 10 days. Percentage of surviving colonies was normalized to the untreated control. Mean ± s.e.m. of n = 4 independent experiments with Student’s t test. **(E)** Proposed model of activation and inhibition of PARP1^V762^. Upon activation, PARP1^V762^ exhibit unfolding of HD helices and new WGR-HD interactions. Furthermore, talazoparib binding at the ART domain drives additional unfolding of the PARP1^V762^ HD, stabilizing interdomain contacts and thereby converting talazoparib into a Type-I PARPi for PARP1^V762^ variant. (F) Allelic distribution and effect of talazoparib on both PARP1 variants indicated on the world map. (G) Restriction fragment length polymorphism at a variant specific cutting site codon 762 and a universal cutting site upstream. PCR fragments were amplified from genomic DNA from different cell lines and digested with EciI then electrophorized on agarose gel.

Next, we assessed the impact of the two PARP1 variants on DNA repair. PARP1 serves a well-established roles as single-strand DNA break (SSB) sensor, recognizing and getting activated by SSBs. The resultant burst of PARylation leads to the recruitment of XRCC1, which is an indispensable scaffold protein in SSB repair, facilitating and accelerating the loading and stabilization of multiple enzymatic components^49^. Probing through immunofluorescence following micro-irradiation, we found that PARPi markedly attenuated the PAR-dependent recruitment of XRCC1, as previously reported^49^ (**Figure 4B**). Notably, there was a prolonged enrichment of XRCC1 at micro-irradiated sites after 30 minutes post-irradiation in PARP1^V762^ bearing cells in the presence of talazoparib, compared to PARP1^A762^ bearing cells (**Figure 4C**). This correlated with the higher retention of PARP1^V762^ at the same sites (**Figure S2B**). Our result suggests that one direct functional consequence of higher PARP1 retention induced by the PARPi talazoparib in the PARP1^V762^ variant might be a stronger entrapment of XRCC1, which would be predicted to affect the resolution of SSBs.

Given the persistent XRCC1 recruitment and accumulated DNA damage in the PARP1^V762^ bearing cells in the presence of talazoparib, we next examined whether there were any differences on the sensitivity of human cells expressing the two PARP1 variants in the presence of the PARPi talazoparib. Using a 10-day colony formation assay titrated with talazoparib, we found that PARP1^V762^ bearing cells exhibited a stronger, dose-dependent sensitivity to talazoparib compared to PARP1^A762^ bearing cells (**Figure 4D**). These data suggest that a link between enhanced PARP1 retention, DNA damage accumulation and better PARPi efficacy of talazoparib for the PARP1^V762^ variant compared to the less frequent but better studied PARP1^A762^ variant.

## DISCUSSION

### Outlook and PARP1 SNP genotyping

Historically nearly all *in vitro* characterization and drug design with PARP1 has been conducted with the less frequent PARP1^A762^ variant and not with the PARP1^V762^ variant, including the mechanistic research on the allosteric activation of the PARP1 enzyme and the dissection of the reverse allostery elicited by distinct PARPi ^5,29^. We surmised that PARP1^V762^ and PARP1^A762^ variants might have a distinct effect on the structure-function relationships owing to the proximity of the PARP1 residue 762 to the PARPi binding site of talazoparib, for example (**Figure S1)**. Through laser micro-irradiation recruitment assay in live cells and using HXMS *in vitro*, we found talazoparib induced a stronger retention of PARP1^V762^ instead of PARP1^A762^, shifting from Type-II (mild allosteric pro-retention for PARP1^A762^) to Type-I PARPi (allosteric pro-retention for PARP1^V762^). While it exerts only modest effects on the HD subdomain and does not alter DNA-binding affinity in PARP1^A762^, as previously reported, it induces reverse allostery in PARP1^V762^, characterized by HD subdomain destabilization and enhanced interdomain contacts (**Figure 4E**). Using the established immunofluorescence-based PARP1 trapping assay, we observed that talazoparib indeed induced stronger trapping of PARP1^V762^ compared to PARP1^A762^ at methylated lesions and further found that this was associated with accumulated DNA damage. Additionally, we observed the persistent recruitment of the single-strand break scaffold protein XRCC1 in PARP1^V762^ bearing cells compared to PARP1^A762^, consistent with augmented DNA damage and the observed hypersensitivity to talazoparib.

Our work suggests there is the need to determine of efficacy and safety differences of PARPi in individuals with different PARP1 variants. Since most i*n vitro* assays and presumably medicinal chemistry campaigns also for PARP1-selective inhibitors may have been conducted with PARP1^A762^, yet the majority of human patients may bear a V762 allele (depending on racial background), we may need to re-assess how this SNP contributes to efficacy and safety in patients across the glove. Our data shows that talazoparib acts like a Type-I PARPi for PARP1^V762^, while functioning as a Type-II PARPi for PARP1^A762^. PARP1^V762^ and PARP1^A762^ variants are all very strongly represented in the human population, but their frequency varies greatly in individuals of Asian ancestry compared to those of African ancestry (**Figure 4F**). Patients with PARP1^V762^ background may thus be more responsive to (and show improved efficacy) a lower dosage of talazoparib, reflected an improved therapeutic index, and/or suffer from higher-grade hematopoietic effects, such as anemia, or gastrointestinal issues including nausea. On the other hand, patients with PARP1^A762^, more common in those with Asian ancestry, might not respond as well to talazoparib treatment.

In order to facilitate further clinical retrospective and prospective work and analysis on the questions which PARP1 alleles are present in a given patient tumor sample, we started to develop a potential diagnostic tool to distinguish the distinct PARP1 alleles. Restriction enzyme EciI recognizes and cuts specially at PARP1^A762^ but not PARP1^V762^ (**Figure 4G**). Along with an upstream universal cutting site, PCR fragments from genomic DNA of U2OS and DLD1 cells clearly showed a cutting pattern for homozygous V762, while those from THP1 and Jurkat cells had double cutting bands and suggested heterozygous alleles. This diagnostic tool and further refinements, such as RT-qPCR based SNP allele mapping in tumors will enable a comprehensive screen of whether there is variant-specific (homozygous A762, homozygous V762, and heterozygous alleles) response to distinct PARPi in the clinic, including progression-free survival, hematological toxicity, GI side-effects, overall tolerability but fundamentally also overall survival, a key criterion over which several PARPi have recently been withdrawn from the cancer clinic.

## Supporting information

HXMS tables

## ACKNOWLEDGMENTS

We thank our colleagues at the University of Pennsylvania, Jennine Dawicki-McKenna and Leland Mayne, for discussions on HXMS experiments. We thank Charlotte Blessing for preliminary life-cell recruitment data with the two PARP1 SNP variants, Tanner Wright for PARP1 enzyme assays, Maren Heimhalt for structural analysis of inhibitor-bound PARP1 variants, as well as Claudia Gonzalez-Leal, Julia Preißer and Andreas Wegerer for project assistance and technical advice. We thank all members of the Ladurner and Margulies teams at LMU Munich for support and discussion, as well as Adrian Schomburg (Eisbach Bio GmbH), Timothy Yap and Guang Peng (both from the MD Anderson Cancer Center) for discussion of PARP1 variants and the clinical use of PARPi. We thank John Pascal and Marie-France Langelier (both from the Université de Montréal) for plasmids. **Funding:** This work was supported by NIH grant CA259037 (to B.E.B) and an Early Career Award from Basser Center for BRCA (to R.B.) and the Deutsche Forschungsgemeinschaft (German Research Foundation; DFG; Project-IDs 213249687 – SFB1064 and 325871075 – SFB1309) (to A.G.L.) and the Ludwig-Maximilians-University of Munich (to A.G.L.). The content is solely the responsibility of the authors and does not necessarily represent the official views of the National Institutes of Health. This manuscript is the result of funding in part by the National Institutes of Health (NIH). It is subject to the NIH Public Access Policy. Through acceptance of this federal funding, NIH has been given a right to make this manuscript publicly available in PubMed Central upon the Official Date of Publication, as defined by NIH.

## Data and materials availability

The HXMS data in this study has been deposited to the PRIDE database (accession code: PXD059341). They will be made public upon publication.

The materials used in this study are available from commercial sources or from the corresponding authors on reasonable request.

## AUTHOR CONTRIBUTIONS

Conceptualization, J.C., R.B., B.E.B. and A.G.L.; methodology, J.C. and R.B.; Investigation, J.C. and R.B.; writing—original draft, J.C. and R.B.; writing—review & editing, B.E.B. and A.G.L.; funding acquisition, B.E.B. and A.G.L.; resources, M.S.C.; supervision, B.E.B. and A.G.L.

## METHODS

### Expression constructs and mutagenesis

pET28 vectors with an N-terminal hexahistidine tag were used to express PARP1^A762^ and PARP1^V762^ (residues 1 to 1014). They were kindly provided by J. Pascal and M.-F. Langelier (U. Montréal).

### Protein expression and purification

Both PARP1^A762^ and PARP1^V762^ were expressed in Rosetta 2 (DE3) *E. coli* and purified using Ni^2+^-affinity, heparin, and gel filtration chromatography^5,42^.

### Hydrogen/deuterium exchange mass spectrometry (HXMS)

2.6 mM of PARP1^A762^ or PARP1^V762^ was incubated for 30 min with 5 mM of DNA containing gap DNA (5’ GCT GGC TTC GTA AGA AGC CAG CTC GCG GTC AGC TTG CTG ACC GCG 3’)^42,46^. To the PARP1/DNA complex, 5.2 mM of talazoparib or olaparib was added and incubated for another 30 min. At room temperature deuterium on-exchange was performed in triplicate (n=3) by adding 15 mL of deuterium on-exchange buffer (10 mM HEPES, pD 7.0, 150 mM NaCl, in D_2_O, pD = pH+0. 4138) to 5 mL of PARP1/DNA/PARPi complex to yield a final D_2_O concentration of 75%. At 100 s, 20 mL aliquots were to 30 mL of ice-cold quench buffer (1.66 M guanidine hydrochloride, 10% glycerol, and 0.8 % formic acid) to make a final pH of 2.4-2.5 and immediately frozen in liquid nitrogen and stored at −80°C. The Non-deuterated (ND) samples were prepared in 10mM HEPES (pH 7.0), 150mM NaCl buffer containing H_2_O and each 20 μL aliquot was quenched into 30 μL of quench buffer. To mimic the on-exchange experiment, the fully deuterated (FD) samples were incubated for 48 h in 75% deuterium but denatured under acidic conditions (0.5% formic acid). 48 h incubation ensures that every amide proton along the entire polypeptide chain undergoes full exchange. Pepsin (Sigma) was coupled to POROS 20 AL support (Applied Biosystems) and the immobilized pepsin was packed into a 64 μL column (2mm x 2 cm, Upchurch). Before electrospraying the samples into the Exactive Plus EMR-Orbitrap (Thermo Fisher Scientific), samples were melted at 0°C and immediately injected into pepsin column and simultaneously pumped at initial flow rate of 50 μL min^-1^ for 2 min followed by 150 μL min^-1^ for another 2 min. The pepsin-digested peptides were trapped onto a TARGA C8 5 μm Piccolo HPLC column (1.0 Å∼ 5.0 mm, Higgins Analytical) and eluted through an analytical C18 HPLC column (0.3 Å∼ 75 mm, Agilent) with a 12–100% buffer B gradient at 6 μL/min (Buffer A: 0.1% formic acid; Buffer B: 0.1% formic acid, 99.9% acetonitrile). MS data acquisition over the mass range 200−2000 m/z were acquired at 60,000 resolution. The effluent was electrosprayed with ion spray voltage of 3.5 kV and capillary temperature operated at 250 °C.

### PARP1 peptide identification

The ND samples were injected into LTQ orbitrap XL, (Thermo Fisher Scientific) for tandem mass spectroscopy (MS/MS). Scan range for MS/MS data was 200–2000 m/z at 15,000 resolution, where the ions were fragmented by CID with normalized collision energy 35. The potential PARP1 peptides were identified through SEQUEST (Bioworks v3.3.1) with a peptide tolerance of 8 ppm and a fragment tolerance of 0.1 AMU against an extensive decoy sequence database (custom database) containing the sequence of both PARP1^A762^ and PARP1^V762^, pepsin and other common contaminants identified in prior HXMS studies. Since pepsin was employed as the protease, we specified it in our search as a non-specific digestion enzyme. After the MS/MS was performed for the first ND, a MATLAB-based program (9.2.0.556344), ExMS2 (version 2017-07-19)^50^ was used with a Ppep score of 0.1 to identify the PARP1 peptides and to generate an exclusion list. While performing MS/MS on the second ND the instrument employed the above exclusion list to collect the MS2 scan of the less intense peptides that were not identified in the previous ND sample. At least four such exclusion lists were generated to increase the number of unique peptides and sequence coverage of the protein. Finally, a pool of peptides based on SEQUEST output files were prepared by EXMS2.

### HXMS analysis and plotting

Peptide pool information generated through ExMS2 was used to identify the deuterated peptides for every sample of HXMS experiment. Each individual deuterated peptide is corrected for loss of deuterium label (backexchange after quench) during HXMS data collection by normalizing to the maximal deuteration level of that peptide in the fully deuterated samples. For each peptide, we have compared the extent of deuteration as measured in the FD sample to the theoretical maximal deuteration (i.e., if no back-exchange occurs). The median extent of back-exchange in our experiments was 24% (**Figure S4A**). The data analysis statistics for all the protein states are in HXMS Supplementary Spreadsheets^51^. The quality of each peptide was further assessed by manually checking mass spectra. The mass spectrometry proteomics data have been deposited to the ProteomeXchange Consortium via the PRIDE^52^ partner repository with the dataset identifier PXD059341. The difference plot for the deuteration levels between any two samples was obtained through an in-house script written in MATLAB. The script compares the deuteration levels between two samples (e.g., PARP1 complex and PARP1/DNA complex) and plots the percent difference of each peptide, by subtracting the percent deuteration of PARP1/DNA complex from PARP1 complex and plotting according to the color legend in stepwise increments (**Figure S3**). The plot of representative peptide data is shown as the mean of three independent measurements +/- SD. Statistical analysis included a t-test with a P < 0.05 (**Figures 1** and **S3**).

### Generation and culture of PARP1-GFP V/A762 cell lines

U2OS ΔPARP1 cells as previously characterized^53^ were transfected with PARP1-GFP-N1 V/A762 plasmids with XtremeGENE HP DNA transfection reagent (Sigma) according to manufacturer’s instructions and selected with 500 μg/mL G418 (Sigma). PARP1-GFP-N1 A762 plasmid has been previously described^53^ and PARP1-GFP-N1 V762 plasmid was generated by site-directed mutagenesis on PARP1-GFP-N1 A762 plasmid with forward primer 5’-CAGTGTGCAGGCCAAGGTGGAAATGCTTGACAACCTGC-3’ and reverse primer 5’-GCAGGTTGTCAAGCATTTCCACCTTGGCCTGCACACTG-3’. The survived cells were further mono-colonized in a 96-well plate at a concentration of 1 cell per well under G418 selection. After two weeks, the visible single colonies were expanded and screened by western blot to choose a comparably expressing pair of variant clones. All U2OS cell lines were cultured in DMEM medium (1 g/L glucose; GIBCO) supplemented with 10% FBS (GIBCO) and 1% penicillin/streptomycin (GIBCO). DLD1, THP1 and Jurkat cell lines were cultured in RPMI-1640 medium (GIBCO) supplemented with 10% FBS and 1% penicillin/streptomycin. Cells were kept inside the cell incubator at an atmosphere of 37 °C and 5% CO_2_ and routinely checked for mycoplasma by PCR-based testing.

### Live-cell imaging following micro-irradiation

Cells were plated in 8-well Nunc Lab-Tek chambers (Thermo Fisher Scientific). Prior to irradiation, cells were treated 1 hour with or without 1 μM of the indicated PARPi (Selleckchem) in Leibovitz’s L-15 medium (GIBCO) supplemented with 10% FBS, at 37 °C in the absence of CO_2_. Live-cell imaging was performed on a Zeiss AxioObserver Z1 confocal spinning-disk microscope with a sCMOS ORCA Flash 4.0 camera (Hamamatsu) and a C-Apo 63x water immersion objective lens. DNA damage was induced along a line of 88 pixels using 10% laser power of a 355 nm laser with 20-22 runs operated through a single-point scanning head (UGA-42 firefly, Rapp OptoElectronics). The recruitment of GFP-tagged proteins at micro-irradiated sites was recorded for 15 min at an interval of 5 s (3 frames pre-damage and first 5 min) and 10 s (last 10 min) and quantified using a custom macro in Fiji/ImageJ^54,55^. The damage region-of-interest (ROI) was selected through a selection of 25 x 100 pixels, while the nucleus ROI was obtained by thresholding the entire nucleus. A background ROI was defined as a region outside of the cell. The recruitment was calculated with the mean fluorescent intensity of each ROI as follows: [Damage(t)-Background(t)]/[Nucleus(t)-Background(t)], relative to the pre-damage value.

### Immunofluorescence

For PARP1 trapping assay, cells were seeded in 96 well SCREENSTAR microplates (Greiner) and treated with PARPi and MMS (Sigma) as indicated for 4 hours the next day. Pre-extraction and fixation were performed as described^47^. In brief, cells were first pre-extracted with a cold cytoskeleton buffer (10 mM PIPES pH 6.8, 300 mM sucrose, 200 mM NaCl, 3 mM MgCl_2_ and 0.6% Triton X-100) for 10 minutes at 4 °C and subsequently fixed with ice-cold methanol for 15 minutes at −20 °C. Cells were then blocked with PBS + 0.1% Tween 20 + 3% BSA for 1 hour at room temperature and incubated with the mouse anti-γH2AX antibody (Millipore, 1:1000 diluted in 1% BSA) overnight at 4 °C. Unspecific antibody staining was removed by washing the cells 3 times in 1% BSA. Subsequently, cells were stained with donkey-anti-mouse Alexa Fluor 555-conjugated fluorescent secondary antibodies (Thermo Fisher, diluted 1:1000 in 1% BSA) along with Hoechst 33342 (Fisher Scientific, 1:5000 diluted) for 1 hour at room temperature. Finally, cells were washed 3 times in 1% BSA and put in PBS while imaging through a Plan-Apochromat 40x/0.95 Korr air objective lens. The fluorescence intensity was quantified by a custom-made ImageJ macro that measures the mean fluorescent signal in the nucleus, based on thresholding cell nuclei using the Hoechst signal.

For XRCC1 recruitment after micro-irradiation, cells were prepared and micro-irradiated as for live-cell imaging. After 5 minutes and 30 minutes post-irradiation, cells were fixed and stained as described^56^. In brief, cells were fixed with 2% paraformaldehyde + 0.1% Triton X-100 for 15 minutes at room temperature, washed 2 times with PBS + 0.1% Triton X-100 and subsequently permeabilized with PBS + 0.1% Triton X-100 for 2 times of 10 minutes at room temperature. After blocking the cells in PBS+ (PBS + 0.5% BSA + 0.15% glycine) for 30 minutes at room temperature, cells were incubated with the mouse anti-XRCC1 antibody (Abcam, 1:1000 diluted in PBS+) overnight at 4 °C. Unspecific antibody staining was removed by washing the cells 4 times in PBS+. Subsequently, cells were stained with donkey-anti-mouse Alexa Fluor 647-conjugated fluorescent secondary antibodies (Thermo Fisher, diluted 1:1000 in PBS+) along with Hoechst 33342 (1:2000 diluted) for 1 hour at room temperature. Finally, cells were washed 4 times with PBS+ and put in PBS while imaging. The fluorescence intensity was analyzed by ImageJ, measuring the mean fluorescence intensity in the micro-irradiated areas and in the complete nucleus.

### Clonogenic survival assay

Cells were seeded in triplicates at a density of 1000 cells per well in 6 well plates and treated continuously with the indicated concentrations of PARP inhibitors from the next day. After 10 days of growth, colonies were fixed and stained with 50% methanol, 7% acetic acid and 0.1% Brilliant Blue R. The colonies were scanned and analyzed with the ColonyArea ImageJ plugin^57^ using a manual threshold of 180. The colony area was used to assess the number of surviving colonies.

### Western blotting

Cells were incubated in lysis buffer (10 mM Tris pH 7.5, 150 mM NaCl, 0.5 mM EDTA, 0.1% SDS, 1% Triton X-100, 1% deoxycholate, and 2.5 mM MgCl_2_) supplemented with complete protein inhibitors cocktail (Roche) for 10 minutes on ice and quantified with the Pierce BCA Protein Assay Kit (Thermo Fisher) and boiled with the same amount of protein for all groups. Samples were then separated with a 4%-15% pre-cast gel (Bio-Rad) and blotted for 25 minutes at 15 V, 1.5 A using a TransBlot Turbo transfer system (Bio-Rad). Blots were blocked in 5% milk in PBS-T (PBS + 0.05% Tween 20) and stained with primary antibodies (rabbit-anti-PARP1 and goat-anti-GFP, homemade; mouse-anti-α-tubulin, Sigma; 1:1,000 diluted in PBS-T) overnight at 4 °C. Blots were washed 2 times of 15 min with PBS-T and incubated with secondary antibodies (Bio-Rad; 1:10,000 diluted in PBS-T) for 1 hour at room temperature. Blots were then washed 3 times with PBS-T and developed using a Curix 60 developer (AGFA).

### Restriction Fragment Length Polymorphism (RFLP) analysis

Cells were harvested and extracted for genomic DNA with ReliaPrep™ gDNA tissue miniprep system (Promega). PCR fragments flanking the codon 762 were amplified by Phusion^™^ high-fidelity DNA polymerase (NEB) with forward primer 5’-TCTCCTCAGATCGACCTTCA-3’ and reverse primer 5’-AAGCTGAGGTGCTGGCTT-3’. PCR products were further loaded onto agarose gel purified through mi-Gel Extraction Kit (metabion). 250 ng of purified products were incubated with 1 μL of EciI (NEB) at 37 °C for 1 hour then inactivated at 65 °C for 20 minutes. Digested products were electrophorized on agarose gel and imaged through Quntum CX5 (Vilber).

### Statistical analyses

All experiments were performed in at least 3 independent biological replicates and compared with Student’s t tests using Prism 8 (Graphpad Software). Significance levels were defined as followed: * p < 0.05, ** p < 0.01, *** p < 0.001, **** p < 0.0001.

## SUPPLEMENTAL FIGURE LEGENDS

**Figure S1.**
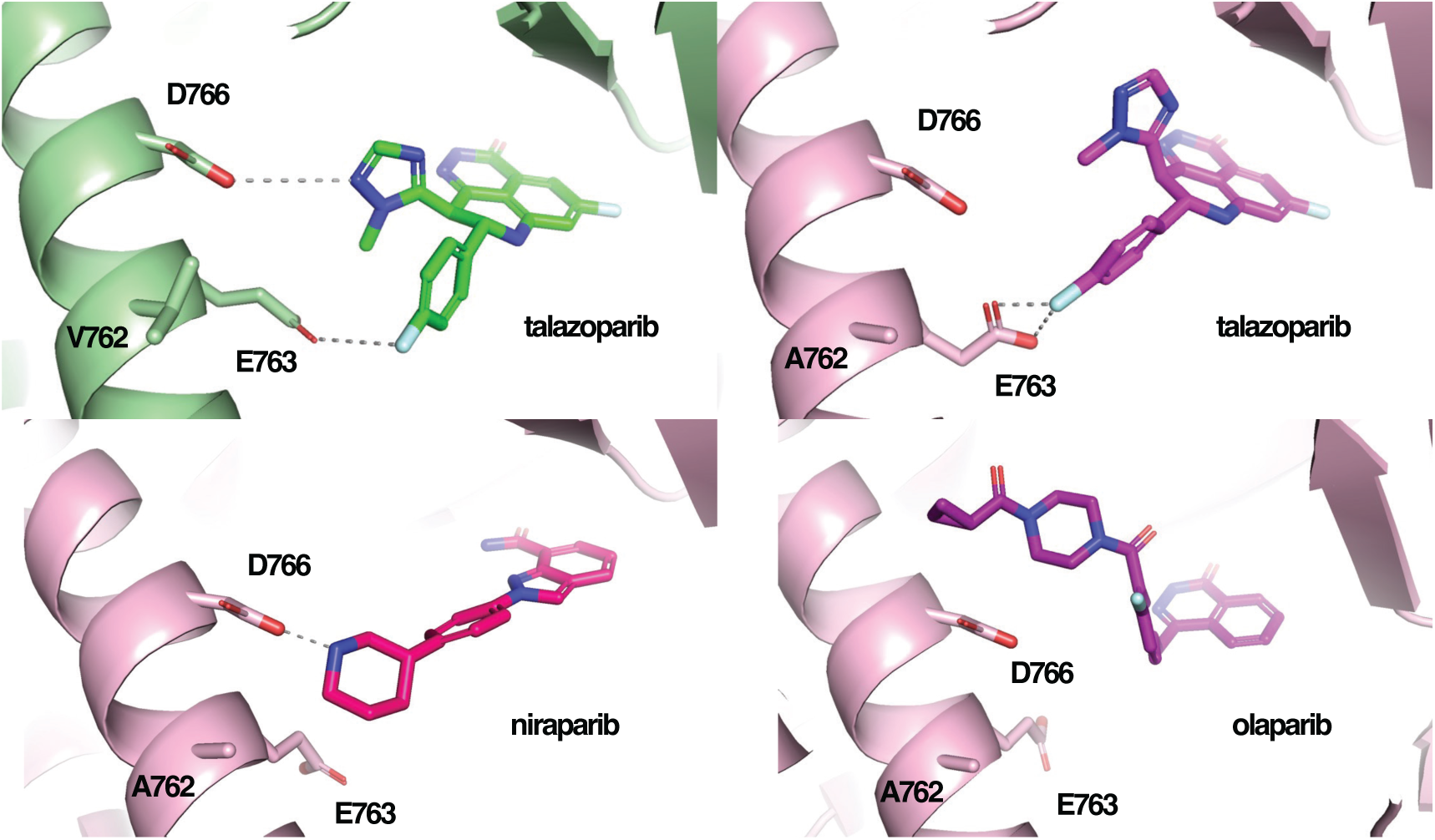
PARP1 variants impact talazoparib binding. Structural alignment of talazoparib-bound PARP1^V762^ CAT (green, PDB: 4PJT) and PARP1^A762^ CAT (magenta, PDB: 4UND) as well as zoom-in of the similar region of niraparib-bound PARP1^A762^ CAT (PDB: 4R6E) and olaparib-bound PARP1^A762^ CAT (PDB: 7AAD).

**Figure S2.**
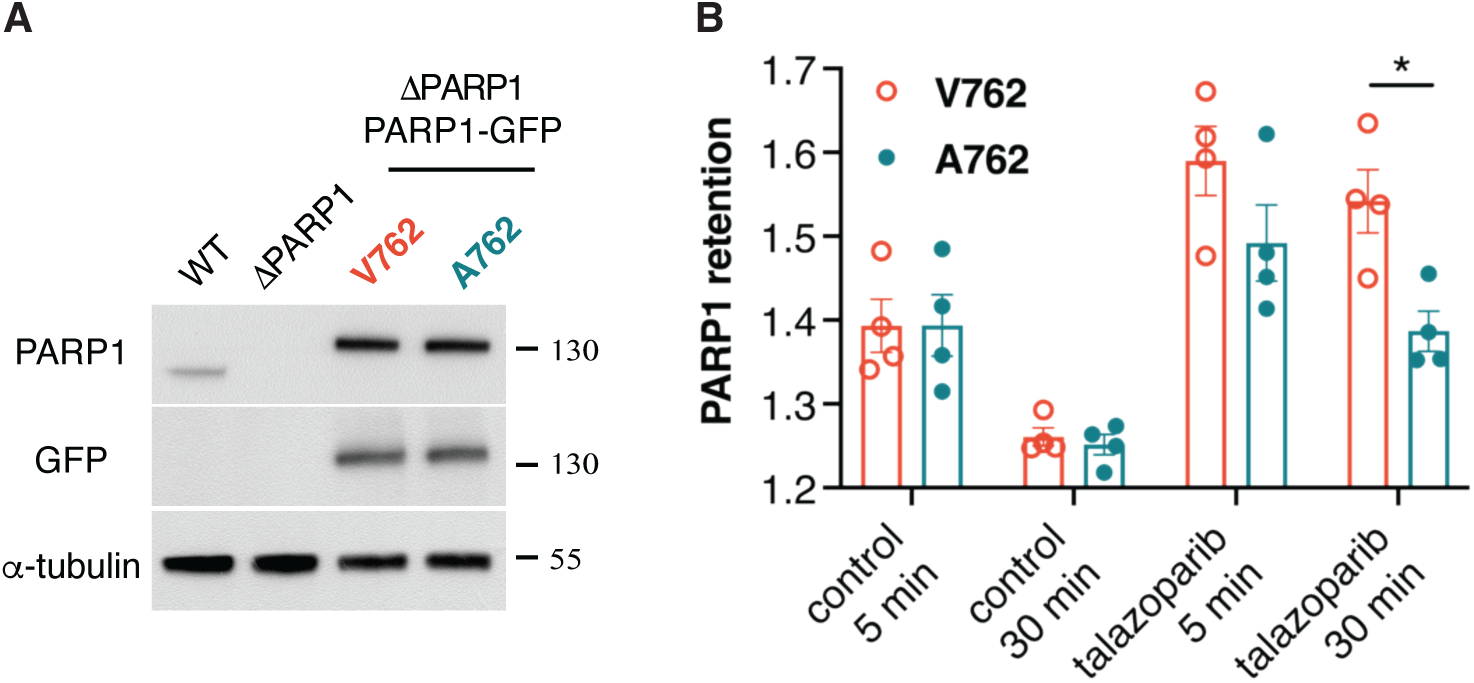
PARP1 variants show comparable expression. (**A**) PARP1 expression in U2OS wild-type (WT) cells, ΔPARP1 cells, and ΔPARP1 cells stably expressing GFP-tagged PARP1^V762^ or PARP1^A762^. Endogenous PARP1 and exogenous PARP1-GFP proteins were probed by anti-PARP1 and anti-GFP antibodies. α-Tubulin was adopted as the loading control. (**B**) Relative recruitment of GFP-tagged PARP1^V762^ or PARP1^A762^ by immunofluorescence after micro-irradiation in U2OS ΔPARP1 cells stably expressing GFP-tagged PARP1^V762^ or PARP1^A762^. Cells were treated with or without 1 μM of talazoparib before irradiation and fixed 30 minutes post irradiation. Mean fluorescence intensity of PARP1 variants at micro-irradiated sites was subtracted to the background intensity and normalized to the nuclear signal. Mean ± s.e.m. of n = 4 independent experiments with Student’s t test.

**Figure S3.**
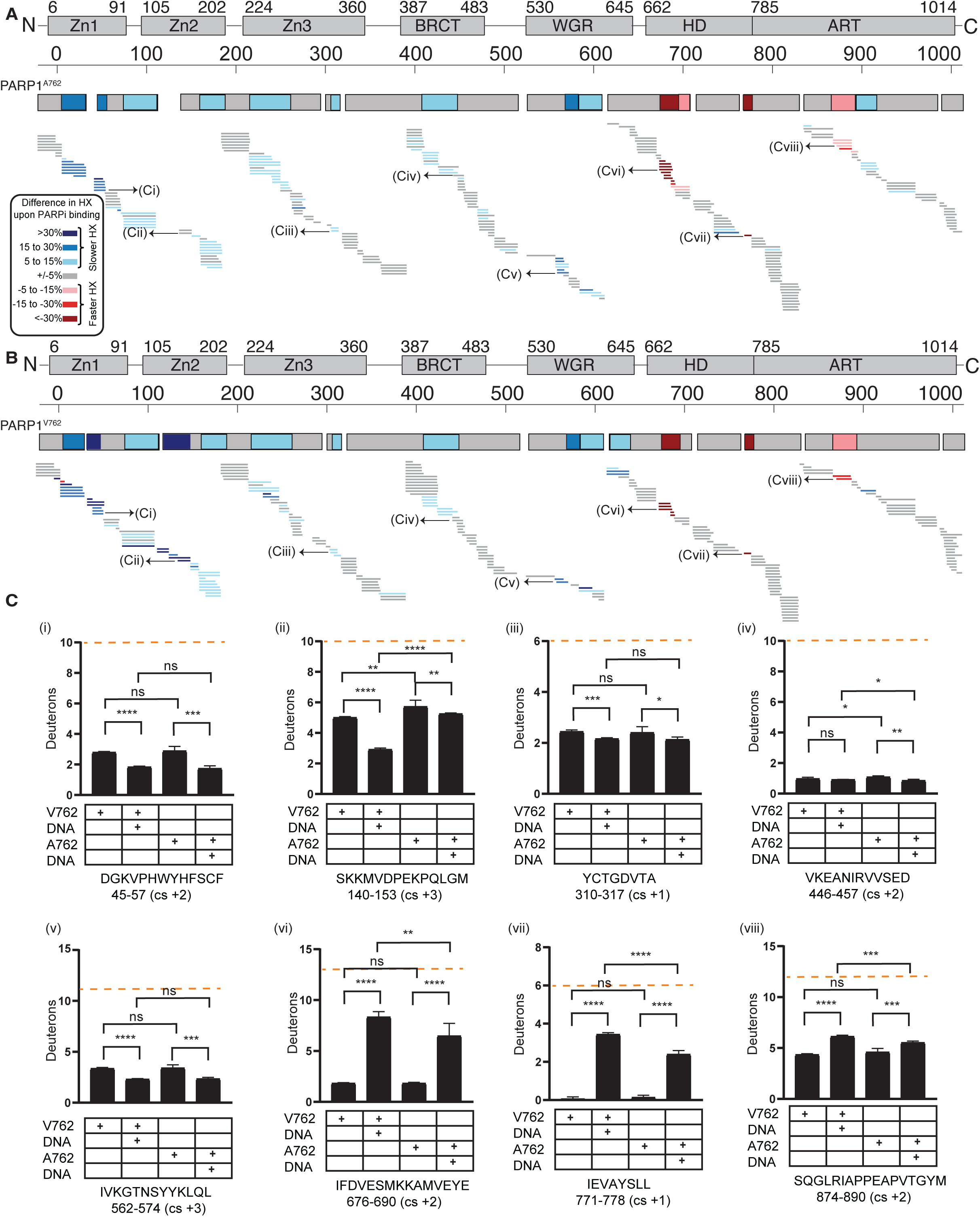
DNA-damage dependent activation of PARP1^V762^. (**A**) The percentage difference in deuteration between PARP1^A762^ and PARP1^A762^ complexed to DNA at 100 s was shown as a HXMS difference plot. Every horizontal bar represents a peptide. The peptides that are in gray represent no change in percentage difference. Peptides in blue represent increased protection and the peptides in red represent decreased protection. The consensus HX difference is shown on the top. The white gaps indicate no peptide coverage. (**B**) The percentage difference in deuteration between PARP1^V762^ and PARP1^V762^ complexed to DNA at 100 s was shown as a HXMS difference plot. Every horizontal bar represents a peptide. The peptides that are in gray represent no change in percentage difference. Peptides in blue represent increased protection and the peptides in red represent decreased protection. The consensus HX difference is shown on the top. The white gaps indicate no peptide coverage. Color key for binning of HX difference is shown at the left corner. (**C**) HX of representative peptides from Zn1, Zn2, Zn3, BRCT, WGR, HD, and ART (i-viii) for PARP1^A762^, PARP1^A762^ complexed with DNA, PARP1^V762^, and PARP1^V762^ complexed to DNA are shown here. An average from three replicates with error bars representing SD and asterisks indicating *P* < 0.05 from two-sided t-test between any two protein states is shown. Maximum number of deuterons (maxD, which is the total number of residues minus the first two residues due to back-exchange within the experimental timescale and minus the number of prolines due to no backbone amide hydrogen) exchanged by the representative peptides is indicated (orange dotted line).

**Figure S4.**
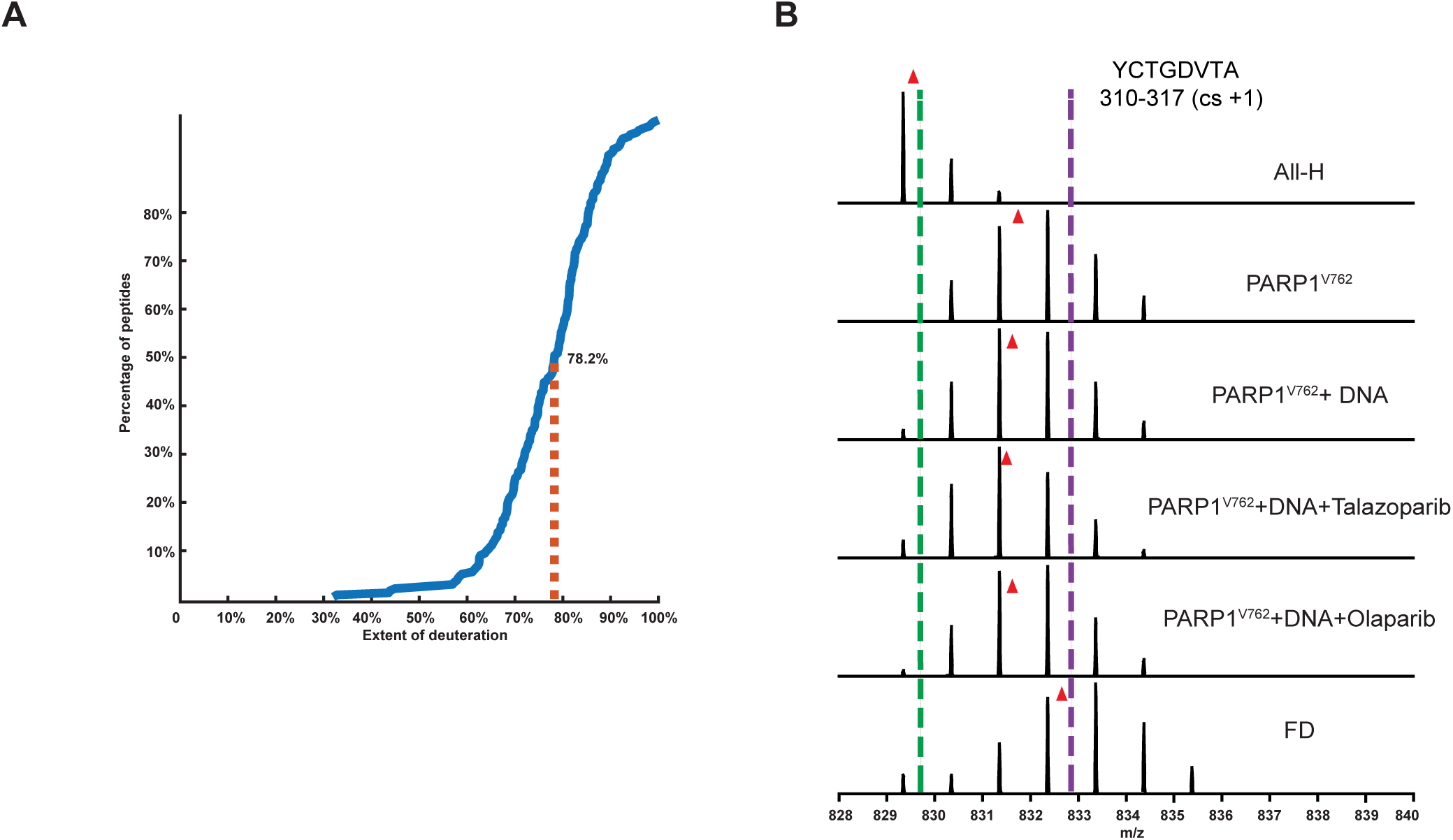
PARP1^V762^ converts talazoparib to an allosteric, pro-retention Type-I PARPi. (**A**) The extent of deuteration is shown as the distribution curve of a representative FD (fully-deuterated) sample. (**B**) Raw MS of representative peptide from Zn3 in All-H (non-deuterated) sample, PARP1^V762^, PARP1^V762^+DNA, PARP1^V762^+DNA+talazoparib, PARP1^V762^ + DNA + olaparib, and the FD sample. Red triangles indicate the centroid value of each protein state. Green and purple dotted lines visualize the differences in m/z of the representative peptide from Zn3.

## REFERENCES

1. Bryant, H.E., Schultz, N., Thomas, H.D., Parker, K.M., Flower, D., Lopez, E., Kyle, S., Meuth, M., Curtin, N.J., and Helleday, T. (2005). Specific killing of BRCA2-deficient tumours with inhibitors of poly(ADP-ribose) polymerase. Nature 434, 913–917. 10.1038/nature03443.

2. Farmer, H., McCabe, N., Lord, C.J., Tutt, A.N.J., Johnson, D.A., Richardson, T.B., Santarosa, M., Dillon, K.J., Hickson, I., Knights, C., et al. (2005). Targeting the DNA repair defect in BRCA mutant cells as a therapeutic strategy. Nature 434, 917–921. 10.1038/nature03445.

3. Amé, J.-C., Rolli, V., Schreiber, V., Niedergang, C., Apiou, F., Decker, P., Muller, S., Höger, T., Murcia, J.M., and De Murcia, G. (1999). PARP-2, A Novel Mammalian DNA Damage-dependent Poly(ADP-ribose) Polymerase. Journal of Biological Chemistry 274, 17860–17868. 10.1074/jbc.274.25.17860.

4. Langelier, M.-F., Planck, J.L., Roy, S., and Pascal, J.M. (2012). Structural Basis for DNA Damage–Dependent Poly(ADP-ribosyl)ation by Human PARP-1. Science 336, 728–732. 10.1126/science.1216338.

5. Dawicki-McKenna, J.M., Langelier, M.-F., DeNizio, J.E., Riccio, A.A., Cao, C.D., Karch, K.R., McCauley, M., Steffen, J.D., Black, B.E., and Pascal, J.M. (2015). PARP-1 Activation Requires Local Unfolding of an Autoinhibitory Domain. Mol Cell 60, 755–768. 10.1016/j.molcel.2015.10.013.

6. Alemasova, E.E., and Lavrik, O.I. (2019). Poly(ADP-ribosyl)ation by PARP1: reaction mechanism and regulatory proteins. Nucleic Acids Research 47, 3811–3827. 10.1093/nar/gkz120.

7. Juarez-Salinas, H., Sims, J.L., and Jacobson, M.K. (1979). Poly(ADP-ribose) levels in carcinogen-treated cells. Nature 282, 740–741. 10.1038/282740a0.

8. Durkacz, B.W., Omidiji, O., Gray, D.A., and Shall, S. (1980). (ADP-ribose)n participates in DNA excision repair. Nature 283, 593–596. 10.1038/283593a0.

9. Satoh, M.S., and Lindahl, T. (1992). Role of poly(ADP-ribose) formation in DNA repair. Nature 356, 356–358. 10.1038/356356a0.

10. Dantzer, F., De La Rubia, G., Ménissier-de Murcia, J., Hostomsky, Z., De Murcia, G., and Schreiber, V. (2000). Base Excision Repair Is Impaired in Mammalian Cells Lacking Poly(ADP-ribose) Polymerase-1. Biochemistry 39, 7559–7569. 10.1021/bi0003442.

11. Murai, J., Huang, S.N., Das, B.B., Renaud, A., Zhang, Y., Doroshow, J.H., Ji, J., Takeda, S., and Pommier, Y. (2012). Trapping of PARP1 and PARP2 by Clinical PARP Inhibitors. Cancer Research 72, 5588–5599. 10.1158/0008-5472.CAN-12-2753.

12. Lord, C.J., and Ashworth, A. (2017). PARP inhibitors: Synthetic lethality in the clinic. Science 355, 1152–1158. 10.1126/science.aam7344.

13. Curtin, N.J., and Szabo, C. (2020). Poly(ADP-ribose) polymerase inhibition: past, present and future. Nat Rev Drug Discov 19, 711–736. 10.1038/s41573-020-0076-6.

14. Drew, Y., Zenke, F.T., and Curtin, N.J. (2025). DNA damage response inhibitors in cancer therapy: lessons from the past, current status and future implications. Nat Rev Drug Discov 24, 19–39. 10.1038/s41573-024-01060-w.

15. Mirza, M.R., Monk, B.J., Herrstedt, J., Oza, A.M., Mahner, S., Redondo, A., Fabbro, M., Ledermann, J.A., Lorusso, D., Vergote, I., et al. (2016). Niraparib Maintenance Therapy in Platinum-Sensitive, Recurrent Ovarian Cancer. N Engl J Med 375, 2154–2164. 10.1056/NEJMoa1611310.

16. Coleman, R.L., Oza, A.M., Lorusso, D., Aghajanian, C., Oaknin, A., Dean, A., Colombo, N., Weberpals, J.I., Clamp, A., Scambia, G., et al. (2017). Rucaparib maintenance treatment for recurrent ovarian carcinoma after response to platinum therapy (ARIEL3): a randomised, double-blind, placebo-controlled, phase 3 trial. The Lancet 390, 1949–1961. 10.1016/S0140-6736(17)32440-6.

17. Robson, M., Im, S.-A., Senkus, E., Xu, B., Domchek, S.M., Masuda, N., Delaloge, S., Li, W., Tung, N., Armstrong, A., et al. (2017). Olaparib for Metastatic Breast Cancer in Patients with a Germline *BRCA* Mutation. N Engl J Med 377, 523–533. 10.1056/NEJMoa1706450.

18. Poveda, A., Floquet, A., Ledermann, J.A., Asher, R., Penson, R.T., Oza, A.M., Korach, J., Huzarski, T., Pignata, S., Friedlander, M., et al. (2021). Olaparib tablets as maintenance therapy in patients with platinum-sensitive relapsed ovarian cancer and a BRCA1/2 mutation (SOLO2/ENGOT-Ov21): a final analysis of a double-blind, randomised, placebo-controlled, phase 3 trial. The Lancet Oncology 22, 620–631. 10.1016/S1470-2045(21)00073-5.

19. Litton, J.K., Rugo, H.S., Ettl, J., Hurvitz, S.A., Gonçalves, A., Lee, K.-H., Fehrenbacher, L., Yerushalmi, R., Mina, L.A., Martin, M., et al. (2018). Talazoparib in Patients with Advanced Breast Cancer and a Germline *BRCA* Mutation. N Engl J Med 379, 753–763. 10.1056/NEJMoa1802905.

20. Golan, T., Hammel, P., Reni, M., Van Cutsem, E., Macarulla, T., Hall, M.J., Park, J.-O., Hochhauser, D., Arnold, D., Oh, D.-Y., et al. (2019). Maintenance Olaparib for Germline *BRCA*-Mutated Metastatic Pancreatic Cancer. N Engl J Med 381, 317–327. 10.1056/NEJMoa1903387.

21. González-Martín, A., Pothuri, B., Vergote, I., DePont Christensen, R., Graybill, W., Mirza, M.R., McCormick, C., Lorusso, D., Hoskins, P., Freyer, G., et al. (2019). Niraparib in Patients with Newly Diagnosed Advanced Ovarian Cancer. N Engl J Med 381, 2391–2402. 10.1056/NEJMoa1910962.

22. Colombo, N., Moore, K., Scambia, G., Oaknin, A., Friedlander, M., Lisyanskaya, A., Floquet, A., Leary, A., Sonke, G.S., Gourley, C., et al. (2021). Tolerability of maintenance olaparib in newly diagnosed patients with advanced ovarian cancer and a BRCA mutation in the randomized phase III SOLO1 trial. Gynecologic Oncology 163, 41–49. 10.1016/j.ygyno.2021.07.016.

23. Kristeleit, R., Lisyanskaya, A., Fedenko, A., Dvorkin, M., De Melo, A.C., Shparyk, Y., Rakhmatullina, I., Bondarenko, I., Colombo, N., Svintsitskiy, V., et al. (2022). Rucaparib versus standard-of-care chemotherapy in patients with relapsed ovarian cancer and a deleterious BRCA1 or BRCA2 mutation (ARIEL4): an international, open-label, randomised, phase 3 trial. The Lancet Oncology 23, 465–478. 10.1016/S1470-2045(22)00122-X.

24. Monk, B.J., Parkinson, C., Lim, M.C., O’Malley, D.M., Oaknin, A., Wilson, M.K., Coleman, R.L., Lorusso, D., Bessette, P., Ghamande, S., et al. (2022). A Randomized, Phase III Trial to Evaluate Rucaparib Monotherapy as Maintenance Treatment in Patients With Newly Diagnosed Ovarian Cancer (ATHENA–MONO/GOG-3020/ENGOT-ov45). JCO 40, 3952–3964. 10.1200/JCO.22.01003.

25. Murthy, P., and Muggia, F. (2019). PARP inhibitors: clinical development, emerging differences, and the current therapeutic issues. CDR. 10.20517/cdr.2019.002.

26. Jain, A., Agostini, L.C., McCarthy, G.A., Chand, S.N., Ramirez, A., Nevler, A., Cozzitorto, J., Schultz, C.W., Lowder, C.Y., Smith, K.M., et al. (2019). Poly (ADP) Ribose Glycohydrolase Can Be Effectively Targeted in Pancreatic Cancer. Cancer Research 79, 4491–4502. 10.1158/0008-5472.CAN-18-3645.

27. Cadzow, L., Brenneman, J., Tobin, E., Sullivan, P., Nayak, S., Ali, J.A., Shenker, S., Griffith, J., McGuire, M., Grasberger, P., et al. (2024). The USP1 Inhibitor KSQ-4279 Overcomes PARP Inhibitor Resistance in Homologous Recombination–Deficient Tumors. Cancer Research 84, 3419–3434. 10.1158/0008-5472.CAN-24-0293.

28. Blessing, C., Apelt, K., Van Den Heuvel, D., Gonzalez-Leal, C., Rother, M.B., Van Der Woude, M., González-Prieto, R., Yifrach, A., Parnas, A., Shah, R.G., et al. (2022). XPC–PARP complexes engage the chromatin remodeler ALC1 to catalyze global genome DNA damage repair. Nat Commun 13, 4762. 10.1038/s41467-022-31820-4.

29. Zandarashvili, L., Langelier, M.-F., Velagapudi, U.K., Hancock, M.A., Steffen, J.D., Billur, R., Hannan, Z.M., Wicks, A.J., Krastev, D.B., Pettitt, S.J., et al. (2020). Structural basis for allosteric PARP-1 retention on DNA breaks. Science 368. 10.1126/science.aax6367.

30. Slade, D. (2020). PARP and PARG inhibitors in cancer treatment. Genes Dev. 34, 360–394. 10.1101/gad.334516.119.

31. Cottet, F., Blanché, H., Verasdonck, P., Le Gall, I., Schächter, F., Bürkle, A., and Muiras, M.-L. (2000). New polymorphisms in the human poly(ADP-ribose) polymerase-1 coding sequence: lack of association with longevity or with increased cellular poly(ADP-ribosyl)ation capacity. J Mol Med 78, 431–440. 10.1007/s001090000132.

32. Yu, H., Ma, H., Yin, M., and Wei, Q. (2012). Association between *PARP-1* V762A polymorphism and cancer susceptibility: a meta - analysis. Genetic Epidemiology 36, 56–65. 10.1002/gepi.20663.

33. Lockett, K.L., Hall, M.C., Xu, J., Zheng, S.L., Berwick, M., Chuang, S.-C., Clark, P.E., Cramer, S.D., Lohman, K., and Hu, J.J. (2004). The *ADPRT V762A* Genetic Variant Contributes to Prostate Cancer Susceptibility and Deficient Enzyme Function. Cancer Research 64, 6344–6348. 10.1158/0008-5472.CAN-04-0338.

34. Hao, B., Wang, H., Zhou, K., Li, Y., Chen, X., Zhou, G., Zhu, Y., Miao, X., Tan, W., Wei, Q., et al. (2004). Identification of Genetic Variants in Base Excision Repair Pathway and Their Associations with Risk of Esophageal Squamous Cell Carcinoma. Cancer Research 64, 4378–4384. 10.1158/0008-5472.CAN-04-0372.

35. Ye, F., Cheng, Q., Hu, Y., Zhang, J., and Chen, H. (2012). PARP-1 Val762Ala Polymorphism Is Associated with Risk of Cervical Carcinoma. PLoS ONE 7, e37446. 10.1371/journal.pone.0037446.

36. Alanazi, M., Pathan, A.A.K., Arifeen, Z., Shaik, J.P., Alabdulkarim, H.A., Semlali, A., Bazzi, M.D., and Parine, N.R. (2013). Association between PARP-1 V762A Polymorphism and Breast Cancer Susceptibility in Saudi Population. PLoS ONE 8, e85541. 10.1371/journal.pone.0085541.

37. Wang, X.-G., Wang, Z.-Q., Tong, W.-M., and Shen, Y. (2007). PARP1 Val762Ala polymorphism reduces enzymatic activity. Biochemical and Biophysical Research Communications 354, 122–126. 10.1016/j.bbrc.2006.12.162.

38. Beneke, S., Scherr, A.-L., Ponath, V., Popp, O., and Bürkle, A. (2010). Enzyme characteristics of recombinant poly(ADP-ribose) polymerases-1 of rat and human origin mirror the correlation between cellular poly(ADP-ribosyl)ation capacity and species-specific life span. Mechanisms of Ageing and Development 131, 366–369. 10.1016/j.mad.2010.04.003.

39. Rank, L., Veith, S., Gwosch, E.C., Demgenski, J., Ganz, M., Jongmans, M.C., Vogel, C., Fischbach, A., Buerger, S., Fischer, J.M.F., et al. (2016). Analyzing structure–function relationships of artificial and cancer-associated PARP1 variants by reconstituting TALEN-generated HeLa *PARP1* knock-out cells. Nucleic Acids Res, gkw859. 10.1093/nar/gkw859.

40. Blessing, C., Mandemaker, I.K., Gonzalez-Leal, C., Preisser, J., Schomburg, A., and Ladurner, A.G. (2020). The Oncogenic Helicase ALC1 Regulates PARP Inhibitor Potency by Trapping PARP2 at DNA Breaks. Molecular Cell 80, 862–875.e6. 10.1016/j.molcel.2020.10.009.

41. Arnold, M.R., Langelier, M.-F., Gartrell, J., Kirby, I.T., Sanderson, D.J., Bejan, D.S., Šileikytė, J., Sundalam, S.K., Nagarajan, S., Marimuthu, P., et al. (2022). Allosteric regulation of DNA binding and target residence time drive the cytotoxicity of phthalazinone-based PARP-1 inhibitors. Cell Chemical Biology 29, 1694–1708.e10. 10.1016/j.chembiol.2022.11.006.

42. Langelier, M.-F., Billur, R., Sverzhinsky, A., Black, B.E., and Pascal, J.M. (2021). HPF1 dynamically controls the PARP1/2 balance between initiating and elongating ADP-ribose modifications. Nat Commun 12, 6675. 10.1038/s41467-021-27043-8.

43. Krishna, M.M.G., Hoang, L., Lin, Y., and Englander, S.W. (2004). Hydrogen exchange methods to study protein folding. Methods 34, 51–64. 10.1016/j.ymeth.2004.03.005.

44. Aoyagi-Scharber, M., Gardberg, A.S., Yip, B.K., Wang, B., Shen, Y., and Fitzpatrick, P.A. (2014). Structural basis for the inhibition of poly(ADP-ribose) polymerases 1 and 2 by BMN 673, a potent inhibitor derived from dihydropyridophthalazinone. Acta Crystallogr F Struct Biol Commun 70, 1143–1149. 10.1107/S2053230X14015088.

45. Thorsell, A.-G., Ekblad, T., Karlberg, T., Löw, M., Pinto, A.F., Trésaugues, L., Moche, M., Cohen, M.S., and Schüler, H. (2017). Structural Basis for Potency and Promiscuity in Poly(ADP-ribose) Polymerase (PARP) and Tankyrase Inhibitors. J. Med. Chem. 60, 1262–1271. 10.1021/acs.jmedchem.6b00990.

46. Velagapudi, U.K., Rouleau-Turcotte, É., Billur, R., Shao, X., Patil, M., Black, B.E., Pascal, J.M., and Talele, T.T. (2024). Novel modifications of PARP inhibitor veliparib increase PARP1 binding to DNA breaks. Biochemical Journal 481, 437–460. 10.1042/BCJ20230406.

47. Illuzzi, G., Staniszewska, A.D., Gill, S.J., Pike, A., McWilliams, L., Critchlow, S.E., Cronin, A., Fawell, S., Hawthorne, G., Jamal, K., et al. (2022). Preclinical Characterization of AZD5305, A Next-Generation, Highly Selective PARP1 Inhibitor and Trapper. Clinical Cancer Research 28, 4724–4736. 10.1158/1078-0432.CCR-22-0301.

48. Bonner, W.M., Redon, C.E., Dickey, J.S., Nakamura, A.J., Sedelnikova, O.A., Solier, S., and Pommier, Y. (2008). γH2AX and cancer. Nat Rev Cancer 8, 957–967. 10.1038/nrc2523.

49. Hanzlikova, H., Gittens, W., Krejcikova, K., Zeng, Z., and Caldecott, K.W. (2016). Overlapping roles for PARP1 and PARP2 in the recruitment of endogenous XRCC1 and PNKP into oxidized chromatin. Nucleic Acids Res, gkw1246. 10.1093/nar/gkw1246.

50. Kan, Z.-Y., Mayne, L., Sevugan Chetty, P., and Englander, S.W. (2011). ExMS: Data Analysis for HX-MS Experiments. J. Am. Soc. Mass Spectrom. 22, s13361-011-0236–3. 10.1007/s13361-011-0236-3.

51. Masson, G.R., Burke, J.E., Ahn, N.G., Anand, G.S., Borchers, C., Brier, S., Bou-Assaf, G.M., Engen, J.R., Englander, S.W., Faber, J., et al. (2019). Recommendations for performing, interpreting and reporting hydrogen deuterium exchange mass spectrometry (HDX-MS) experiments. Nat Methods 16, 595–602. 10.1038/s41592-019-0459-y.

52. Perez-Riverol, Y., Csordas, A., Bai, J., Bernal-Llinares, M., Hewapathirana, S., Kundu, D.J., Inuganti, A., Griss, J., Mayer, G., Eisenacher, M., et al. (2019). The PRIDE database and related tools and resources in 2019: improving support for quantification data. Nucleic Acids Research 47, D442–D450. 10.1093/nar/gky1106.

53. Sellou, H., Lebeaupin, T., Chapuis, C., Smith, R., Hegele, A., Singh, H.R., Kozlowski, M., Bultmann, S., Ladurner, A.G., Timinszky, G., et al. (2016). The poly(ADP-ribose)-dependent chromatin remodeler Alc1 induces local chromatin relaxation upon DNA damage. MBoC 27, 3791–3799. 10.1091/mbc.E16-05-0269.

54. Rueden, C.T., Schindelin, J., Hiner, M.C., DeZonia, B.E., Walter, A.E., Arena, E.T., and Eliceiri, K.W. (2017). ImageJ2: ImageJ for the next generation of scientific image data. BMC Bioinformatics 18, 529. 10.1186/s12859-017-1934-z.

55. Schindelin, J., Arganda-Carreras, I., Frise, E., Kaynig, V., Longair, M., Pietzsch, T., Preibisch, S., Rueden, C., Saalfeld, S., Schmid, B., et al. (2012). Fiji: an open-source platform for biological-image analysis. Nat Methods 9, 676–682. 10.1038/nmeth.2019.

56. Mandemaker, I.K., Zhou, D., Bruens, S.T., Dekkers, D.H., Verschure, P.J., Edupuganti, R.R., Meshorer, E., Demmers, J.A.A., and Marteijn, J.A. (2020). Histone H1 eviction by the histone chaperone SET reduces cell survival following DNA damage. Journal of Cell Science 133, jcs235473. 10.1242/jcs.235473.

57. Guzmán, C., Bagga, M., Kaur, A., Westermarck, J., and Abankwa, D. (2014). ColonyArea: An ImageJ Plugin to Automatically Quantify Colony Formation in Clonogenic Assays. PLoS ONE 9, e92444. 10.1371/journal.pone.0092444.

